# Subthreshold psychosis symptoms associated with molecular genetic risk in a population-based cohort: Findings from Generation Scotland

**DOI:** 10.1101/821041

**Authors:** A.R. Docherty, Andrey A. Shabalin, Daniel E. Adkins, Frank Mann, Robert F. Krueger, Archie Campbell, Caroline Hayward, David J. Porteous, Andrew M. McIntosh, Kenneth S. Kendler

## Abstract

**Importance:** Subthreshold psychosis symptoms in the general population may be associated with genetic risk for schizophrenia. In this analysis, empirically-derived symptom factor scores led to a detection of significant and robust polygenic signal.

**Objective:** This study sought to optimize genetic association with data-driven symptom factor scores, accounting for cohort factor structure and sex differences.

**Design:** EFA-derived symptom factor scores were regressed onto PRS for schizophrenia in models accounting for age and genetic ancestry principal components. Follow-up examination of symptom factor score associations with other related genetic risks included ADHD, autism, bipolar disorder, major depression, and neuroticism.

**Participants:** This study examined the newly expanded symptom dataset from the Northern European ancestry cohort, Generation Scotland: Scottish Family Health Study (*N* = 9,105 individuals 18-65 years of age) comprising common variant and subthreshold psychosis symptom data. A total of 5,391 females and 3,713 males with age M[SD] = 45.2 [13] were included in the final analyses.

**Main Outcome and Measure:** Subthreshold psychosis symptoms were measured using the *Schizotypal Personality Questionnaire-Brief* (SPQ-B). Primary phenotypic factor scores and genome-wide polygenic risk scores (PRS) reflected weighted sum scores and were examined as continuous measures. Polygenic risk scores were calculated from genome-wide association summary statistics using 7,358,674 imputed common genetic variants passing quality control.

**Results:** In males, symptom factor scores were positively associated with polygenic risk for schizophrenia alone and implicated primarily interpersonal/negative symptoms. In females, symptom factor scores were positively associated with polygenic risks for ADHD and autism but not schizophrenia. Scores were robustly associated with genetic risk for neuroticism across both males and females.

**Conclusions and Relevance:** This study detected a significant association of subthreshold psychosis symptoms with genetic risk for schizophrenia and neuroticism in a population-based sample. Furthermore, important sex differences suggest a need for better understanding of schizophrenia risk assessment in females.

**Key Points:** *Question:* What molecular genetic risks are associated with subthreshold psychosis symptoms in the general population?

*Findings:* In a large population-based cohort (*N* = 9,084), significant associations of polygenic risks with symptoms were observed. Symptoms were associated with genetic risk for schizophrenia in males, for ADHD and autism spectrum disorder in females, and for neuroticism across both males and females.

*Meaning:* Associations of genetic risk with symptoms in the general population are highly significant and suggest important sex differences.

## Introduction

Subthreshold psychosis symptoms, sometimes referred to as schizotypy,^1–7^ have served as the basis for myriad studies of genetic risk for psychosis. Biometrical analyses of these symptoms have produced significant heritability estimates ranging from .3 to .5,^8–10^ with biological relatives typically evincing increased negative/interpersonal symptoms. Subthreshold psychosis symptoms are milder than overt symptoms, are frequently observed in the general population, and can be measured continuously.

However, association of these symptoms with molecular genetic risk for schizophrenia has not yet been detected in a population-based sample. One well-powered molecular genetic analysis has recently detected shared genetic covariance of psychotic experiences (e.g., auditory/visual hallucinations and delusions) with schizophrenia and with four other psychiatric disorders using interview data from UK Biobank.^4^ These results are promising for genetic studies of subthreshold psychosis symptoms, since schizotypy measures provide increased variation and sensitivity with respect to risk—a key empirical finding from the literature on subthreshold negative symptoms.^11–13^

The availability of symptom and common variant genetic data in a large population-based cohort, the Generation Scotland Scottish Family Health Study (*N* = 9,105 individuals aged 18-65 years), yields the unique opportunity to examine associations of these symptoms with polygenic risk for schizophrenia. Because of potential for clinically significant overlap of items with other psychiatric symptoms, we followed up with additional analyses of polygenic risk for the four other psychiatric disorders examined in analyses of UK Biobank—major depression, bipolar disorder, autism spectrum disorder, and ADHD.^4^

Recent data from the World Health Organization have suggested that psychotic experiences may be much less specific to schizophrenia than previously thought.^1^ Indeed, psychotic experiences appear to lie on the continuum of neuroticism, and have been observed to either precede or follow the onset of a range of non-psychotic psychiatric disorders.^14–17^ Thus, it was also expected that symptoms would be associated with genetic risk for neuroticism—consistent with dimensional conceptualizations of psychosis, and with empirical evidence of high shared genetic covariance of psychosis with neuroticism.^18–20^

## Methods

### Sample

Data were comprised of adults ages 18-65 from Generation Scotland: The Scottish Family Health Study (GS; data available on request at http://www.generationscotland.co.uk). GS is a family-and population-based study consisting of 23,690 participants recruited from general medical practices across Scotland. The protocol and sample characteristics are described in detail elsewhere.^21^ Briefly, participants were all adults and were not ascertained on the basis of having any particular disorder. It is possible that individuals with diagnoses of psychosis or a mood disorder were recruited as part of this sample, but the use of a population-based sample was thought to capture a cohort representative of the general population. Sample characteristics for these analyses are presented in Table 1. GS received ethical approval from the NHS Tayside Committee on Medical Research Ethics (REC Reference Number: 05/S1401/89).

**Table 1.**
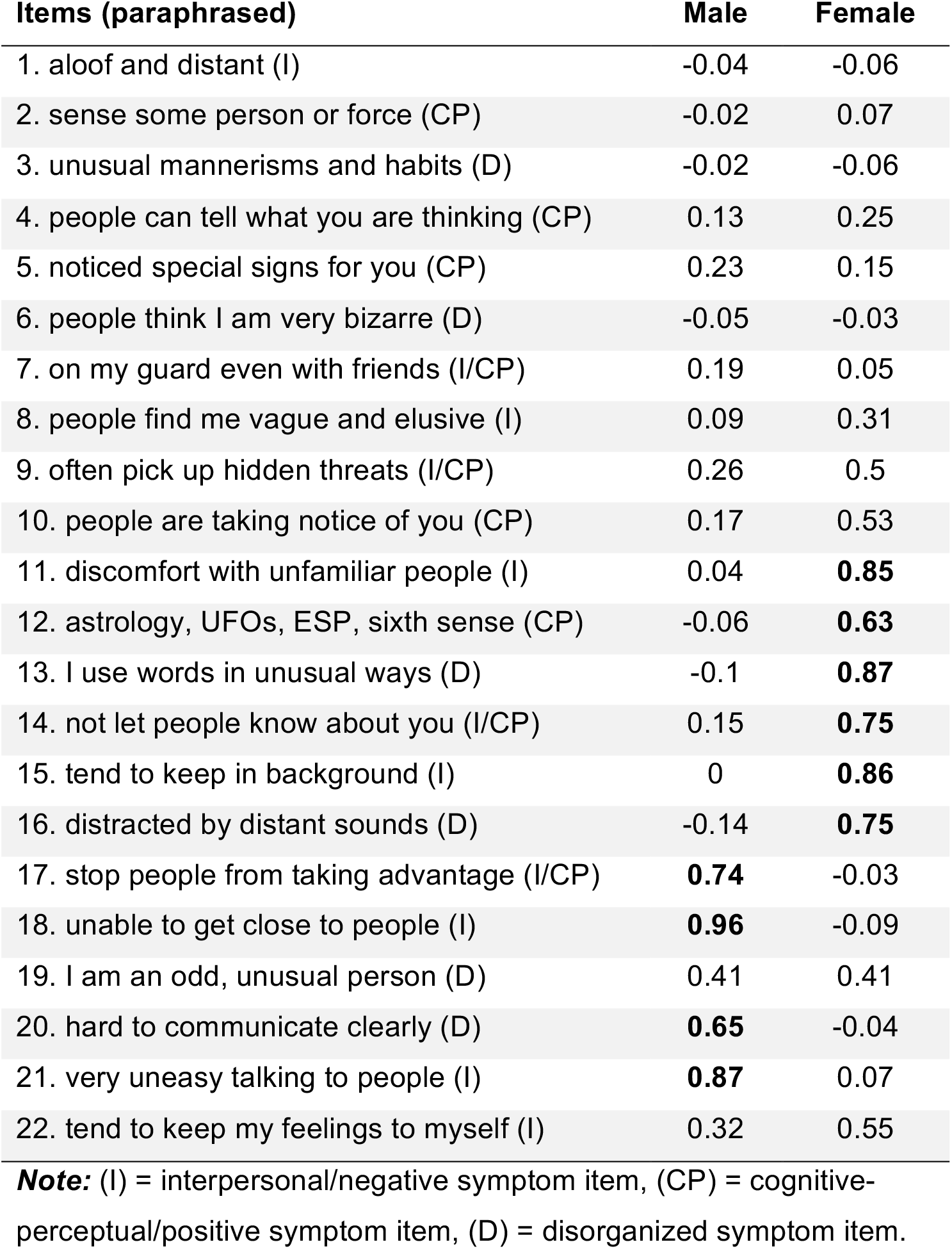
Items and primary factor loadings from male and female exploratory factor analysis models.

### Phenotypic Measurement

The oft-used and well-validated *Schizotypal Personality Questionnaire* (SPQ; Raine 1991) taps subthreshold psychosis symptoms and psychotic-like experiences.^22^ Items are thought to reflect a phenotypic-indicator of liability for schizophrenia-spectrum disorders.^23^ The brief version (the 22-item SPQ-B)^24^ measures a full range of symptoms based on the operational definition of Schizotypal Personality Disorder (SPD).^25^ These items reflect the same self-report information found on interview measures such as the Structured Interview for Schizotypy (SIS)^2^ and the Structured Interview for Prodromal Symptoms (SIPS).^26^ According to the International Consortium on Schizotypy Research and in current common parlance, *schizotypy* and *schizotypal* are now virtually interchangeable given the strong degree of measurement overlap. These symptoms are broadly cognitive (e.g., paranoid ideation, ideas of reference), interpersonal (e.g., anhedonia, no close friends), and behavioral (e.g., odd behavior and language).^27,28^ SPQ-B items have been widely used with schizophrenia probands, their first-degree relatives, adolescents, twins, outpatients, and college students,^3,7,24,29,30^ and the SPQ is the most commonly utilized questionnaire for assessing these symptoms across cultures and languages. Notably, there is considerable variability in total and subscale scores across healthy cohorts (See **Table 1** for the means and variances of SPQ-B total scores in Generation Scotland), but demonstrated reliability and several sources of evidence for validity of the measure.^30,31^

### Factor Structure and Analysis of Measurement Invariance

EFA with oblique rotation, using the parallel analysis fa() function in the nFactors R package, identified three factors in the full cohort. Follow-up confirmatory factor analysis using multiple group nested likelihood ratio tests indicated highly significant measurement noninvariance across sex in tests of metric invariance (loadings Δ*X*^2^ = 744.54, *df* = 60, *p* < 0.001), as well as tests of strict measurement invariance (loadings, intercepts, residual variances (Δ*X*^2^ = 2400.43, *df* = 104, *p* < 0.001). To derive sex-specific factors, EFAs were fit separately in females and males (**Figure S1**) and weighted sum scores were calculated from primary factor item loadings. The use of only the primary symptom factor scores reduced the number of tests run, and the use of factor scores bypassed typical issues of zero-inflation in non-clinical cohorts, common to psychosis-related items. Symptom scores were significantly negatively correlated with age in males (r = −.22, p < .0001) and not in females (r = .01, ns). Symptom scores were not significantly associated with any of the ancestry principal components.

### Genotyping and Imputation

DNA collection and calling for Generation Scotland are detailed elsewhere.^21^ Genotypes were imputed using the Michigan Imputation Server pipeline, including phasing via Eagle^32^ and imputation via minimac3^33^ using 1000 Genomes as a reference panel. SNPs with ambiguous strand orientation, >5% missing calls, or out of Hardy-Weinberg equilibrium (p < 0.001) were excluded. SNPs with minor allele frequency below 0.01 or imputation R^2^ < 0.5 were also excluded. PLINK^34^ was used to perform quality control. Final polygenic risk scoring was performed on 7,358,674 variants passing quality control.

### Polygenic Risk Scoring

Polygenic risk scores were calculated using PRSice 2.0^35^ based on genome-wide association summary statistic weights from the largest current GWAS meta-analyses.^36–40^ Previous studies have utilized multiple p-value thresholds to create PRS with increasing portions of genomic data to detect changes in *R*^2^. To minimize the number of exploratory tests, a default *a priori* p-value threshold of 1.0 was selected using the maximum number of variants available.^41^ Modest but significant differences were observed between males and females in PRS for schizophrenia, with higher PRS for schizophrenia in males (t = 2.53; p = .01). Other differences included ADHD PRS, derived from sex-specific summary scores.

### Regressions of Symptoms onto Polygenic Risk

Generation Scotland evidenced normal distributions of all PRS and positive skew of the symptom factor score in both females and males. Cube transformation was sufficient to correct this skew. No differences in prediction of any item were detected when using mixed models to account for cryptic relatedness (**Figure S2**). Linear regressions of the primary symptom factor onto schizophrenia PRS included age and the first three genetic ancestry principal components as covariates. Separate models in females and males were compared with and without PRS, examining Nagelkerke rsq and the PRS coefficient in the multivariate model. Follow-up tests examined the other PRS in similar model comparisons.

## Results

A total of 5,391 females and 3,713 males were included in the final analyses. **Table 1** presents the EFA-derived primary factor item loadings in females and males. In males, factor scores reflected higher loadings on negative/disorganized symptoms. In females, factor scores reflected a mix of symptoms. Sample characteristics and symptom factor scores for males, females, and the entire cohort are presented in **Table 2**. Symptom scores were significantly positively associated with genetic risk for schizophrenia in males but not in females (males rsq = .0024; *t* = 2.610, *p* = 0.009; females rsq = .0026, t = −2.092, p = .037).

**Table 2.** Demographics, Symptoms, and Empirically Derived Symptom Factor Scores

| **Sample** | **Females** | **Males** | **Entire Cohort** |
| --- | --- | --- | --- |
| *N* | 5391 | 3713 | 9104 |
| Age *M(SD)* | 45.2 (13.3) | 45.2 (13.7) | 45.2 (13.4) |
| SPQ-B Sum Score *M(SD)* | 3.9 (3.7) | 3.9 (3.7) | 3.9 (3.7) |
| Symptom Factor Score *M(SD)* | 1.6 (1.8) | 0.4 (0.9) |  |

*Note:* M = mean, SD = standard deviation, EFA = exploratory factor analysis. Schizotypy factor scores are specific to male and female EFAs.

In follow-up tests, associations in males were specific to schizophrenia risk and were not observed for ADHD, autism, bipolar disorder, or major depression. Symptom scores in females were significantly positively associated with genetic risk for ADHD and for autism spectrum disorders (ADHD rsq = .0064; *t* = 3.268, *p* = .001; autism rsq = .0091; *t* = 3.876, *p* = 1.2×10^−4^) and were not associated with genetic risk for bipolar disorder or major depression. Finally, schizotypy scores accounted for a significant proportion of variance in genetic risk for neuroticism in males (0.5%) and in females (2.1%) (males rsq = .0051; *t* = 3.828, *p* = 1.3×10^−4^; females rsq = .0209; *t* = 9.080, *p* = 3.7×10^−9^).

## Discussion

This study is the first to detect statistically significant association of schizotypy with polygenic risk for schizophrenia in the general population. Results were specific to males, with higher levels of negative symptoms and higher schizophrenia PRS. Associations of symptoms in females with ADHD and autism genetic risk warrants further follow-up with respect to what psychopathology this symptom measure might be tapping in females.

Use of empirically-derived symptom factors based on the test cohort allowed us to detect variation in molecular genetic risk signal specific to schizophrenia in males. In contrast to findings in males, results suggest that items primarily endorsed by females on the SPQ-B are related to genetic risk for autism or ADHD, rather than for Cluster A psychopathology. Based on previous research, there is reason to believe that other PRS may be robust to variation in schizotypy symptoms: PRS for ADHD has predicted attention problems,^42^ PRS for autism has predicted cognitive ability,^43^ and PRS for major depression has predicted variation in antidepressant treatment response^44^ and could be associated with self-reported anhedonia. Indeed, items with the highest loadings in females pertain to some of the common cognitive and interpersonal experiences of individuals with ADHD and autism.

Importantly, the strongest associations observed in this study relate to genetic risk for neuroticism. Recent progress in psychiatric genetics has led to further consolidation and metaanalysis of phenotypic and molecular data to more effectively model the latent structure of the psychosis spectrum and higher-order dimensions of psychopathology risk.^45,46^ We may find that dimensional rather than categorical polygenic risks best predict psychosis, but that is yet to be determined. Enhanced quantitative approaches may further refine what we consider to be psychosis risk, and will ideally involve several methods and measures, multiple genetic risk metrics, and careful attention to psychometrics within each population being measured.^47–50^

### Limitations

One limitation of this study is a reliance on questionnaire rather than interview data. However, the use of such questionnaires will be a prerequisite of any large population study, and the measure has strong overlap with conventional interviews. Another limitation is the lack of a clinical or pre-clinical comparison cohort. Factor analysis of psychotic-like symptoms in diverse psychiatric cohorts could generate different symptom dimensions that may more closely approximate “true” psychosis risk. Another limitation relates to the generalizability of our findings to cohorts of smaller sizes, as effects are small. It is important to note that while statistical associations are robust, the variance accounted for is very modest, and in any given polygenic risk study rarely reaches 3%. Finally, findings may not generalize to other ancestries, despite the global appeal of SPQ-B items, because (and very unfortunately) genome-wide association studies for these PRS still almost exclusively rely on Northern European cohorts. It bears mentioning that with current methods, examination of diverse risk groups enhances GWAS discovery and increases generalizability.^51–53^

### Future Directions

Family and molecular genetic studies have provided evidence that a negative schizophrenia symptom dimension may hold predictive utility^13,54–58^ but phenotyping in genomic studies has been light, or samples too small, to adequately address questions about the common variant genetic architecture of psychosis symptoms in the general population. It appears possible that dimensional conceptualizations of a psychosis continuum that incorporate normative experience will compliment categorical approaches with respect to polygenic risk prediction. Given the observed relationships in this study, we believe it is likely that future genetic risk research (particularly with respect to psychosis and other low base-rate disorders) will benefit from attention to sex differences, measurement noninvariance, cohort factor structure, and perhaps association with polygenic risk for higher-order dimensions of psychopathology (e.g., neuroticism).^59,60^

## Acknowledgements and funding

This project was supported by the National Institute of Mental Health (grant number K01MH093731; A.D.), a Brain & Behavior Research Foundation Young Investigator Award (A.D.), the Simons Foundation (A.D.), the American Foundation for Suicide Prevention (A.D.), and the Tanner Foundation (A.S.). Generation Scotland received core support from the Chief Scientist Office of the Scottish Government Health Directorates [CZD/16/6] and the Scottish Funding Council [HR03006]. Genotyping of the Generation Scotland: Scottish Family Health Study samples was carried out by the Genetics Core Laboratory at the Wellcome Trust Clinical Research Facility, Edinburgh, Scotland and was funded by the Medical Research Council UK and the Wellcome Trust (Wellcome Trust Strategic Award “Stratifying Resilience and Depression Longitudinally” (STRADL) Reference 104036/Z/14/Z).

---

**Supplemental Figure 1.**
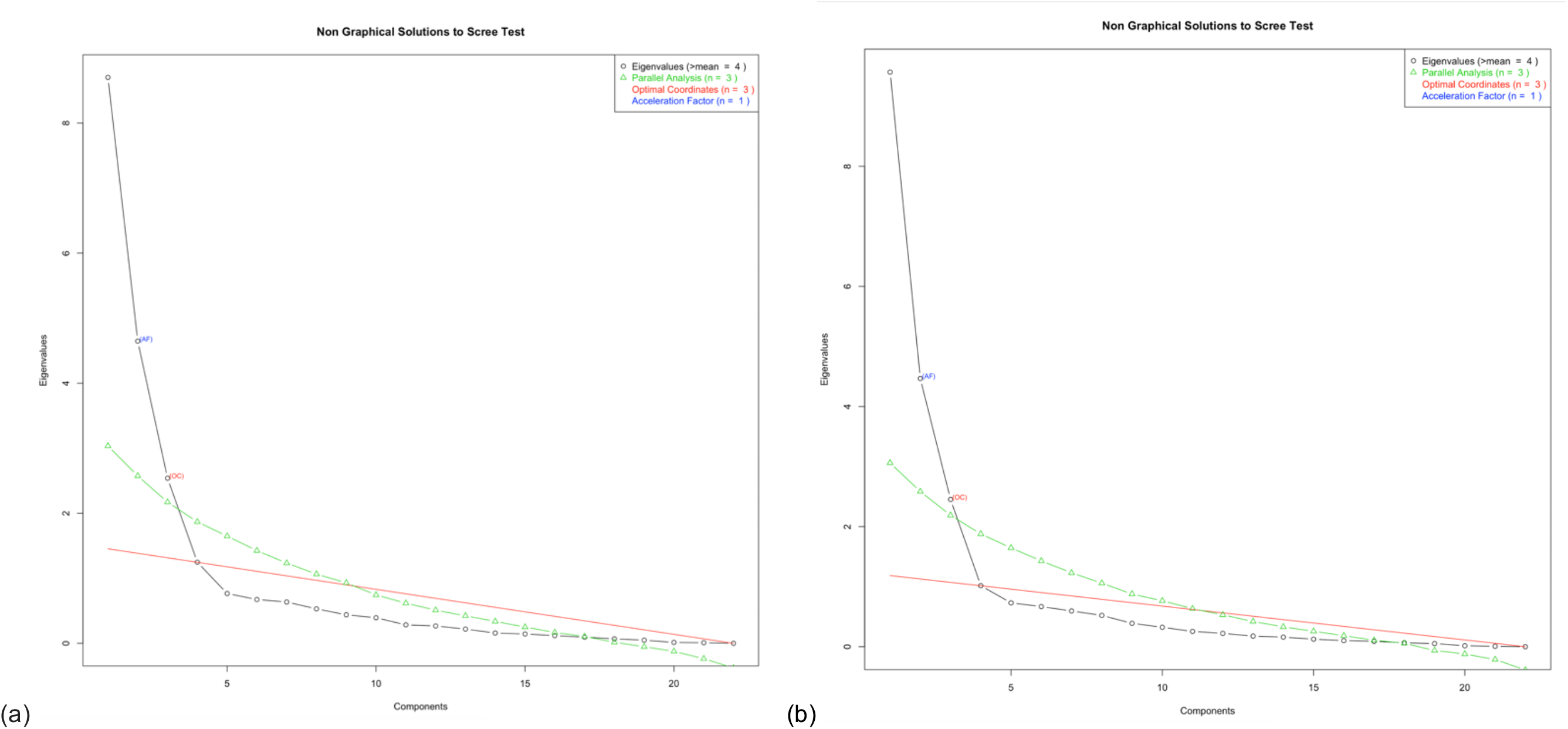
Eigen plots of the factor structure of psychosis proneness symptoms in Generation Scotland males (a) and females (b) point to three factors in both groups.

**Supplemental Figure 2.**
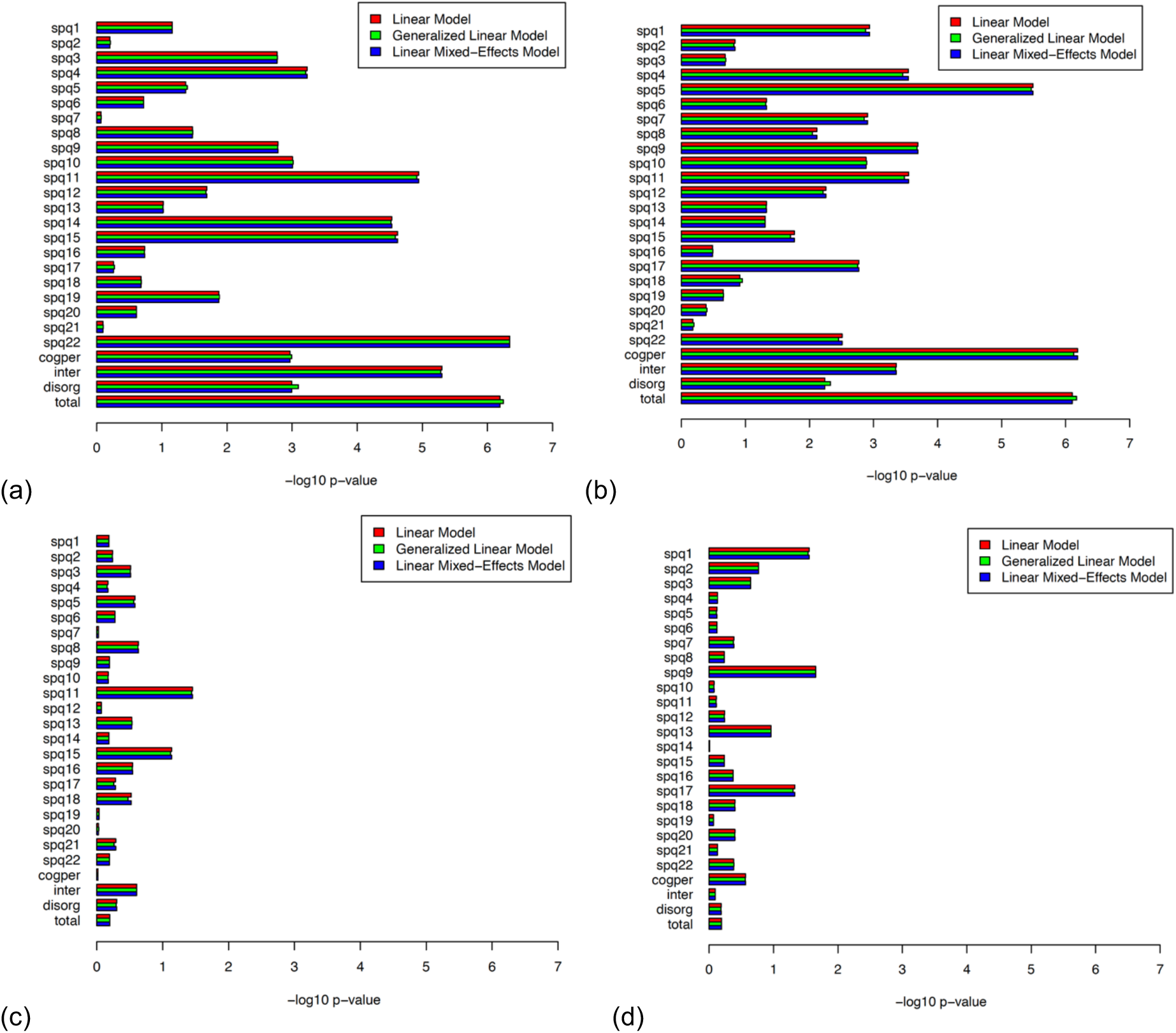
No differences in model fit/prediction were detected when using mixed models to account for cryptic relatedness. Plots (a) and (b) show prediction of individual items with neuroticism PRS in females and males, respectively. Plots (c) and (d) show prediction using schizophrenia PRS.

